# Assessing HLA imputation accuracy in a West African population

**DOI:** 10.1101/2023.01.23.525129

**Authors:** Ruth Nanjala, Mamana Mbiyavanga, Suhaila Hashim, Santie de Villiers, Nicola Mulder

## Abstract

The Human Leukocyte Antigen (HLA) region plays an important role in autoimmune and infectious diseases. HLA is a highly polymorphic region and thus difficult to impute. We therefore sought to evaluate HLA imputation accuracy, specifically in a West African population, since they are understudied and are known to harbor high genetic diversity. The study sets were selected from Gambian individuals within the Gambian Genome Variation Project (GGVP) Whole Genome Sequence datasets. Two different arrays, Illumina Omni 2.5 and Human Hereditary and Health in Africa (H3Africa), were assessed for the appropriateness of their markers, and these were used to test several imputation panels and tools. The reference panels were chosen from the 1000 Genomes dataset (1kg-All), 1000 Genomes African dataset (1kg-Afr), 1000 Genomes Gambian dataset (1kg-Gwd), H3Africa dataset and the HLA Multi-ethnic dataset. HLA-A, HLA-B and HLA-C alleles were imputed using HIBAG, SNP2HLA, CookHLA and Minimac4, and concordance rate was used as an assessment metric. Overall, the best performing tool was found to be HIBAG, with a concordance rate of 0.84, while the best performing reference panel was the H3Africa panel with a concordance rate of 0.62. Minimac4 (0.75) was shown to increase HLA-B allele imputation accuracy compared to HIBAG (0.71), SNP2HLA (0.51) and CookHLA (0.17). The H3Africa and Illumina Omni 2.5 array performances were comparable, showing that genotyping arrays have less influence on HLA imputation in West African populations. The findings show that using a larger population-specific reference panel and the HIBAG tool improves the accuracy of HLA imputation in West African populations.

**Author Summary:** For studies that associate a particular HLA type to a phenotypic trait for instance HIV susceptibility or control, genotype imputation remains the main method for acquiring a larger sample size. Genotype imputation, process of inferring unobserved genotypes, is a statistical technique and thus deals with probabilities. Also, the HLA region is highly variable and therefore difficult to impute. In view of this, it is important to assess HLA imputation accuracy especially in African populations. This is because the African genome has high diversity, and such studies have hardly been conducted in African populations. This work highlights that using HIBAG imputation tool and a larger population-specific reference panel increases HLA imputation accuracy in an African population.

## Introduction

The human Major Histocompatibility Complex (MHC) region, also known as the Human Leukocyte Antigen (HLA) region, is a large locus in the human genome composed of a set of polymorphic genes. It is located on the short arm of the human chromosome 6 with over 200 genes, 128 of which are predicted to be expressed (1). It spans around 5Mbp and contains more than 250,000 Single Nucleotide Polymorphisms (SNPs). It is one of the most complex regions in the human genome because of its high density of polymorphism and linkage disequilibrium. The HLA region is classified into 3 main classes; HLA class I, HLA class II and HLA class III and lies on chromosome 6 between positions 29Mb and 34Mb (2). The HLA region is associated with cancer development (3), the innate and adaptive immune system, cord blood and bone marrow transplants, a wide range of autoimmune and infectious diseases (4), the complement cascade system and adverse drug reactions (5). Identifying the exact genetic variants in the HLA region associated with diseases is of utmost importance to discover the underlying genetic pathophysiology (6) and identify potential therapeutic targets.

To identify the specific alleles that are associated with immune responses and immune-mediated traits, Genome-wide Association Studies (GWAS) use microarrays to genotype these genes at a moderate cost (7). However, these arrays are limited as they can only measure a small number of the SNPs and thus are limited in the number of variants that can be accurately assayed (8).The HLA region is highly variable as the alleles are inherited in a Mendelian fashion from each parent and thus vary from individual to individual (9). This further complicates the collection and use of genotyping data using genotyping arrays in large cohorts. To curb this limitation and dissect the variation of the HLA loci in GWAS, genotype imputation is conducted, taking into consideration the long-range disequilibrium between the HLA loci and SNP markers across the HLA region (10).

Compared to other populations, African genomes are more diverse and have a reduced linkage disequilibrium, making it even more difficult to impute HLA alleles (11). Africa is regarded as the cradle of modern humans, *Homo sapiens. Populations* on other continents descended from groups that migrated from Africa thousands of years ago. It is considered the most genetically diverse continent in the world as African genomes retained more variation than other world populations (12). Genome-wide SNP genotyping study findings indicate that African populations, for example, have maintained a large and subdivided structure throughout evolutionary history (13), and that the deepest splits between human populations lie in Sub-Saharan Africa (14), (15).

Accurately imputing the HLA region is key, considering its role in immune responses. Assessing imputation accuracy is necessary as imputation works based on statistical inferences which involve probabilities. Imputation performance can be affected by several factors such as genotyping arrays, number of individuals in the reference panel, the genetic and ethnic diversity represented, data quality, statistical method of the imputation tools and how well the reference and study panels match.

The aim of the study was to assess the accuracy of imputing the HLA region in a West African population, as HLA imputation accuracy in Africans has not been extensively studied, despite the heaviest disease burden occurring in Africa (16). Previous studies like (17), have focused on assessing general imputation accuracy in African populations rather than HLA imputation accuracy. The few studies that have done HLA imputation accuracy in African populations have used African Americans as the target dataset (18). This study determined the accuracy of HLA imputation with respect to software, reference panel selection, reference panel sample size and genotyping arrays. The performance of 4 imputation tools, three HLA-specific and one general, was tested. In addition, the effect of a population-specific versus a non-population-specific reference panel on imputation in a West African population was tested. Finally, the impact of using data genotyped on different platforms and reference sample sizes for HLA imputation was also assessed.

These results help to inform future GWAS studies on the most appropriate software, recommended reference panel for HLA imputation and the influence of genotyping arrays and reference panel size on HLA imputation accuracy.

## Results

### Sample data

The target dataset was obtained from the GGVP WGS dataset and used to select markers on the H3Africa and Illumina Omni 2.5 arrays. Of the 1,731,033 SNP markers on the H3Africa array, 13,435 matched those in the GGVP WGS dataset, while 1,717,596 were unique to the H3Africa array. Of the 2,314,963 SNP markers on the Illumina Omni 2.5 array, 13,850 SNPs matched those in the GGVP WGS dataset, while 2,301,113 were unique to the Illumina Omni 2.5 array.

### Imputation concordance

Table 1 shows the overall concordance rate for the different imputation tools, genotyping arrays, and reference panels. Compared to HLA typing, the overall concordance rate of the imputed data was 0.84 for HIBAG, 0.77 for Minimac4, 0.58 for SNP2HLA and 0.17 for CookHLA. The HLA Multi-ethnic was the best performing reference panel with an accuracy rate of 0.87, followed by the H3Africa panel at 0.619, then 0.609 for 1kg-Afr, 0.604 for 1kg-All and 0.531 for 1kg-Gwd. For the array comparison, data created from the Omni 2.5 array was more accurate followed by data created from the H3Africa array. The Omni 2.5 array contained a few more Gambian SNPs than the H3Africa array, which would likely impact results.

**Table 1:** Overall Concordance rate

|  | Reference panels |  |  |  |  |  |
| --- | --- | --- | --- | --- | --- | --- |
|  | 1kg-All | 1kg-Afr | 1kg-Gwd | H3Africa | HLA<br>Multi-ethnic |  |
| Imputation<br>program |  |  |  |  |  | Average |
| SNP2HLA | **0.65633** | 0.63917 | 0.425 | 0.61483 | - | 0.58383333 |
| HIBAG | 0.83 | 0.85017 | 0.77883 | **0.88967** | - | 0.83716667 |
| CookHLA | 0.10083 | 0.17217 | 0.20767 | **0.21183** | - | 0.173125 |
| Minimac4 | 0.82717 | 0.7765 | 0.711 | 0.7605 | **0.873** | 0.76879167 |
| **Average** | 0.603583 | 0.6095 | 0.530625 | 0.619208 | 0.873 |  |

In terms of Concordance rate, the H3Africa reference panel outperformed the 1kg-All, 1kg-Afr, and 1kg-Gwd reference panels when using HIBAG (0.89) and CookHLA (0.21) for imputation. For SNP2HLA, 1kg-All (0.66) outperformed 1kg-Afr, 1kg-Gwd and H3Africa. For data imputed using Minimac4, the HLA Multi-ethnic reference panel (0.87) performed better than the other reference panels. There was no comparison of SNP2HLA, HIBAG and CookHLA on the HLA Multi-ethnic panel as the Michigan imputation server that contains the reference panel was prebuilt with Minimac4 only. From the analysis, HLA-C allele imputation was found to be most accurate, followed closely by HLA-A and lastly HLA-B (Table 2).

**Table 2:**
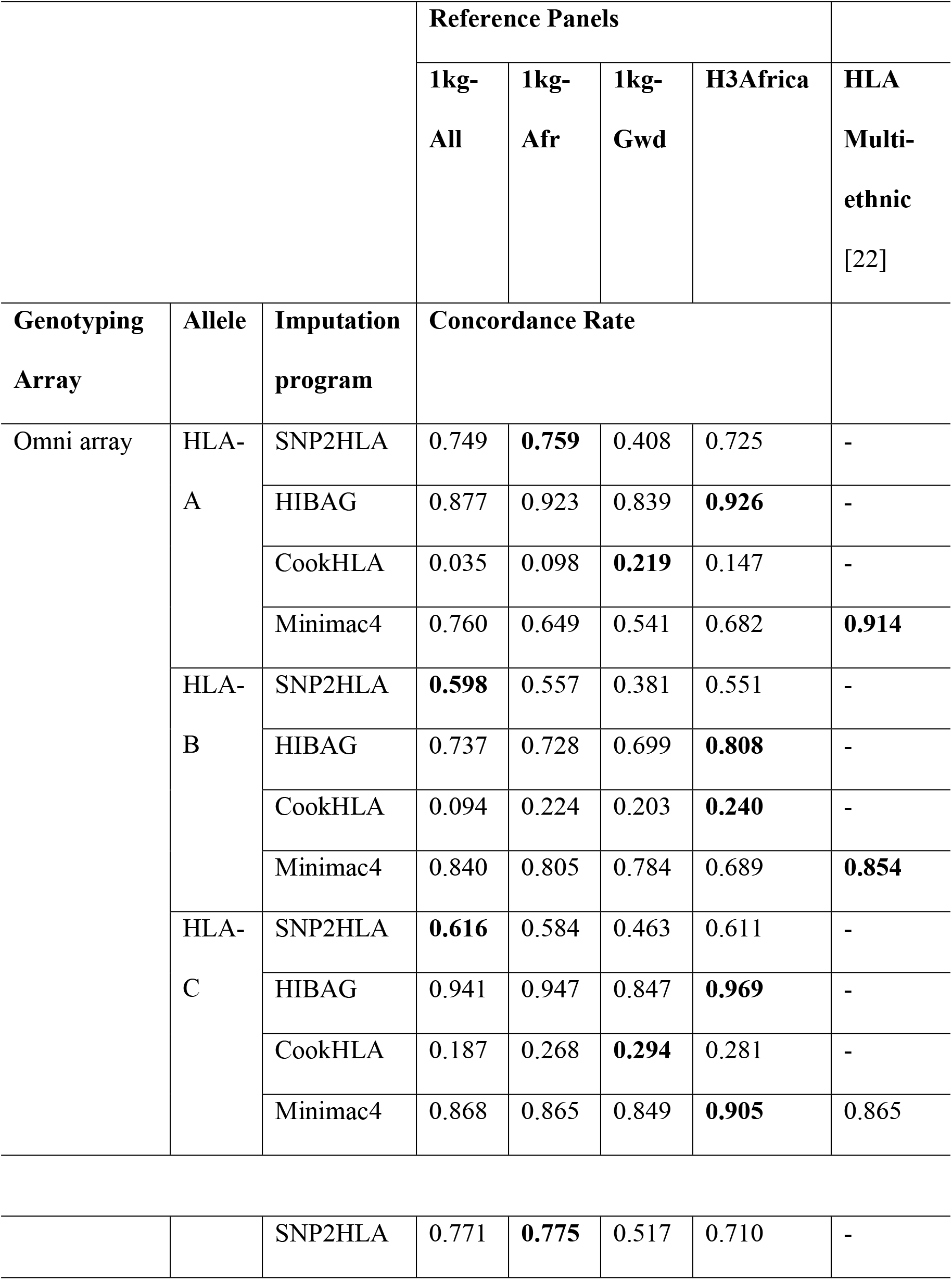
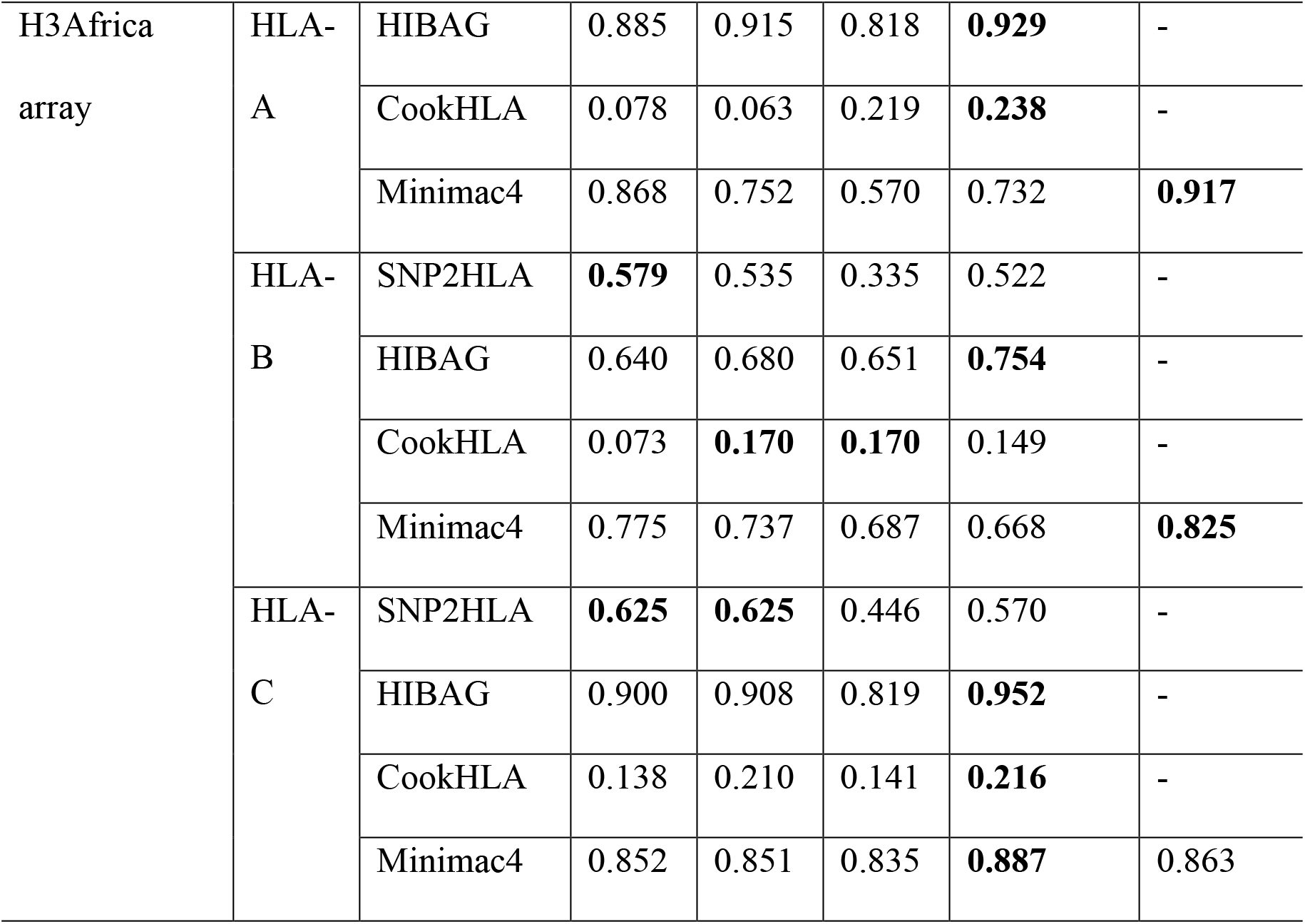
Allele-Specific Concordance rate

### Imputation accuracy based on reference panels

The H3Africa reference panel had the highest concordance with HLA typing when HIBAG (0.89) and CookHLA (0.21) tools were used. For SNP2HLA (0.66), the 1kg-All was the best performing reference panel, while the HLA Multi-ethnic had the highest concordance rate when using Minimac4 (0.87) (Fig 1).

**Fig 1:**
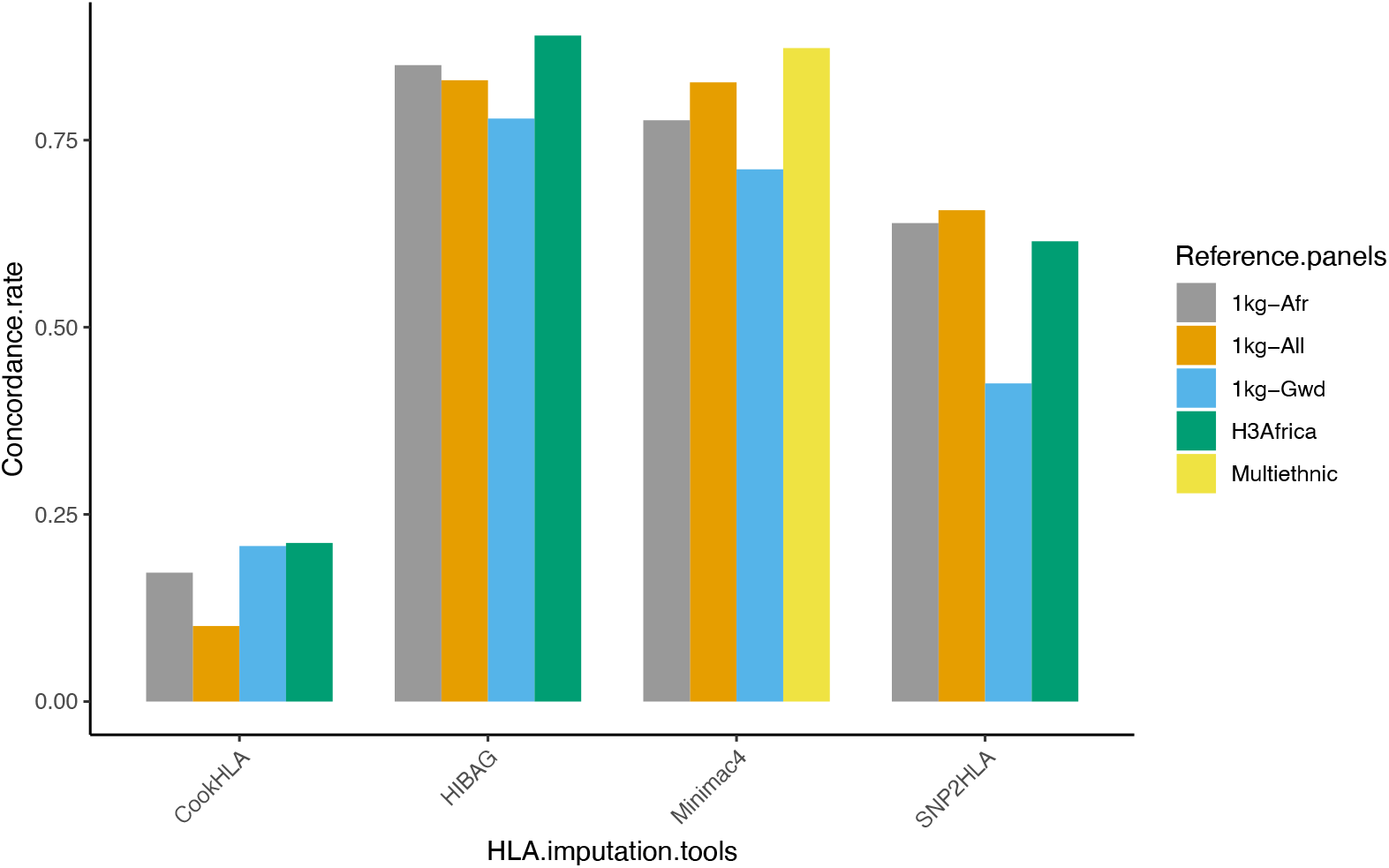
Concordance rate based on reference panels. Generally, the H3Africa reference outperformed the 1kg-All, 1kg-Afr and 1kg-Gwd reference panels.

### Comparison of Allele frequency and accuracy of HIBAG

As HIBAG was the best performing imputation tool, HLA alleles imputed by HIBAG were used for allele frequency and accuracy rate comparison and the output is plotted in Fig 2. HLA imputation accuracy dropped when the frequency of HLA alleles increased across all the reference panels, especially for the HLA-B alleles. This is comparable to a study by (19), who demonstrated that most low frequency HLA alleles had high concordance rates in African Americans and European Americans.

**Fig 2:**
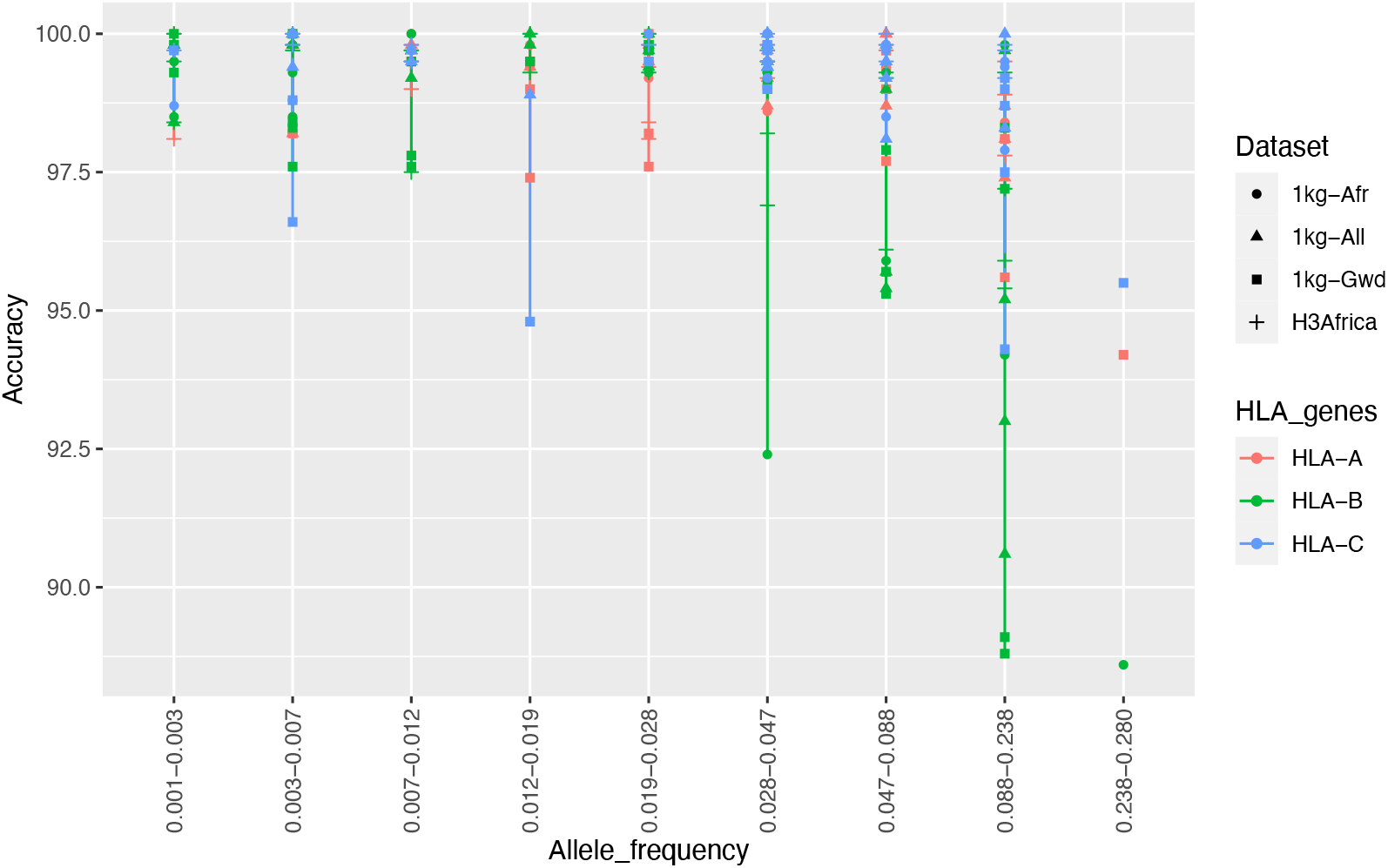
Allele frequency vs Accuracy of HIBAG. Accuracy tended to decrease with increasing frequency, especially for HLA-B alleles.

### Imputation accuracy based on error rates

Overall, HLA-B alleles had higher error rates showing they were imputed less accurately. CookHLA imputed HLA alleles with the highest error rates (Fig 3a). HLA-B seemed to have higher error rates for SNP2HLA and HIBAG, while HLA-A alleles had higher error rates for CookHLA and Minimac4.

**Fig 3:**
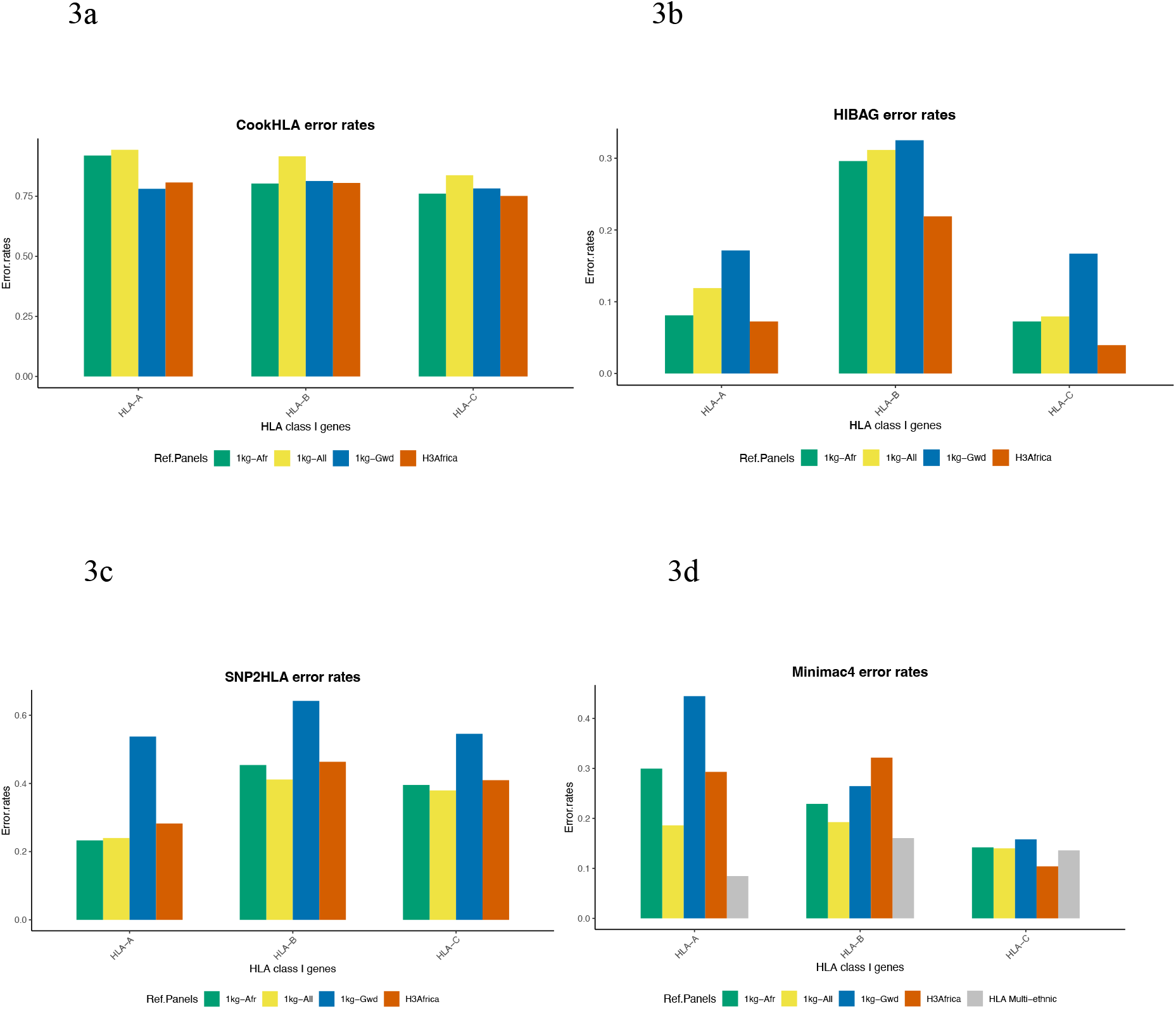
Imputation Accuracy comparison based on error rates. Results from HIBAG (Fig 3b) showed that HLA-B had a higher error rate, followed by HLA-A and lastly, HLA-C. For SNP2HLA (Fig 3c), HLA-B imputation was less accurate, followed by HLA-C and finally HLA-A. HLA-A had higher error rates for Minimac4 (Fig 3d) and CookHLA (Fig 3a), followed by HLA-B and finally HLA-C alleles.

An interesting observation was that Minimac4, a general imputation tool, imputed HLA-B alleles more accurately than any of the HLA-specific imputation tools.

## Discussion

Accurate imputation of classical HLA alleles is key for association studies to understand the genetic risk of autoimmune and infectious diseases. According to (20), imputation of classical HLA alleles offers an invaluable additional layer of interrogation of the variation in the HLA region. Still, it should by no means be expected to fully explain an observed association as a SNP could be tagging an effect in one of the HLA genes.

Assessing HLA imputation accuracy is critical because HLA is a highly variable region, and imputation is a statistical procedure based on probabilities. HLA imputation accuracy has previously been conducted in non-West African populations and admixed populations but not exclusively in West African populations. Therefore, a detailed comparison of five reference panels, four imputation tools, two genotyping arrays, and reference panel sample size in a West African population was provided in this study. The tools were selected based on their proven performance in previous studies.

The study focused on imputing HLA class I genes, that is, HLA-A, HLA-B and HLA-C alleles using SNPs tagging the HLA region. Imputation of HLA-B was less accurate compared to HLA-C and HLA-A imputation because HLA-B alleles are highly polymorphic (21) compared to HLA-A and HLA-C. According to (22), there are over 3000 allelic variants in the HLA-B region. However, accurate imputation of HLA-B alleles is important, as they play a key role in the progression of acquired immune deficiency syndrome. Slow progression of the disease has been associated with individuals expressing HLA-B*57 and HLA-B*27, while rapid progression has been associated with individuals expressing HLA-B*35 alleles (23). Minimac4 showed improved imputation accuracy of HLA-B alleles, suggesting that a general imputation tool can be used for studies targeting HLA-B alleles.

Another important factor is the choice of genotyping array. The Illumina Omni 2.5 array performance was slightly better than that of the H3Africa array because it has more SNPs in the target population, 13,850 SNPs compared to 13,436 SNPs. This difference was, however, statistically insignificant, showing that the choice of genotyping arrays has little influence on HLA imputation accuracy. Note, however, that the two arrays have significant overlap in their content, which may explain the similarities. Therefore, a comparison of more diverse arrays is necessary to fully assess the impact of array content.

In (24), it was shown that genome-wide coverage of genotyping arrays correlates with the number of SNPs on the genotyping arrays but does not correlate with the imputation quality. Therefore, the choice of genotyping arrays should be based on additional genotyping array content such as pharmacogenetics or HLA variants and not only on the extent of genome coverage of genotyping arrays.

HIBAG outperformed Minimac4, SNP2HLA and CookHLA in terms of imputation accuracy. This is because HIBAG is robust for populations with complex linkage disequilibrium blocks (5). Compared to Minimac4, SNP2HLA and CookHLA, HIBAG uses unphased genotyped data, eliminating variation provided by phasing software and shortening the computational phasing steps. In terms of computational burden, HIBAG takes long to run when the reference panel needs to be customized. For instance, the 1kg-All reference panel, which was the largest, took approximately 20 days and 32 threads when training with HIBAG compared to a few hours with 9 threads when training with SNP2HLA. SNP2HLA provides an added advantage over HIBAG as it imputes HLA SNPs, amino acids, and alleles, unlike HIBAG, which imputes only HLA alleles.

Generally, the size of the reference panel has a substantial impact on HLA allele imputation accuracy (25). As expected, increased accuracy was achieved with a more extensive reference panel. To assess the accuracy of imputing HLA alleles using a larger reference panel, we performed imputation on the large HLA Multi-ethnic reference panel via the Michigan imputation server. Imputation accuracy was slightly higher than the other reference panels, but we could not compare it with the other tools as the server only provides the Minimac4 tool. Other than the reference panel sample size, population specificity also affected imputation accuracy.

Overall, the H3Africa reference panel outperformed the other reference panels due to its larger sample size and relatedness to the target population. It outperformed the other panels when imputing using HIBAG and CookHLA, while the 1kg-All reference performed better when imputing using SNP2HLA. This implied that HIBAG’s and CookHLA’s performance was based on population specificity and sample size, while SNP2HLA’s performance was based on sample size alone.

## Conclusion and future recommendations

HLA allele imputation uses the correlation between the HLA genes and nearby SNPs in the reference panel to type unknown HLA alleles from SNP array data in the target dataset. HIBAG, CookHLA, Minimac4 and SNP2HLA imputation tools were used to impute HLA alleles.

The most effective software for HLA allele imputation in West African populations in this study was HIBAG. However, it takes a lot of time and memory during the training of the reference panel. Reference panel sample size and population content influence HLA allele imputation accuracy, which was found to decrease when allele frequency increased.

This study identified factors to consider when selecting an imputation tool and reference panel with respect to informing association studies that focus on the HLA region and West African populations. The results highlight the best tools and panels to use for accurately imputing HLA genotypes.

A recommendation will be to test additional African populations other than the Gambian population as it would better assess imputation accuracy in specific African populations. Reference panels comparable in size are also needed when testing imputation accuracy to reduce biasness in the study, where a single large reference panel outperforms the reference panels that are smaller in size.

Also, there is need to build large African-specific reference panels to enable high-quality imputations for studies that cannot afford the cost of next-generation sequencing. The aim being to generate more data that can be used for genome-wide association and fine mapping studies in African populations.

## Materials and Methods

### Study populations

The reference panels were chosen from the 1000 Genomes dataset (1kg-All), 1000 Genomes African dataset (1kg-Afr), 1000 Genomes Gambian dataset (1kg-Gwd), Human Hereditary and Health in Africa (H3Africa) dataset and the HLA Multi-ethnic dataset. The 1kg-All consists of 26 world populations (26), the 1kg-Afr is the African dataset drawn from the 1kg-All dataset, while the 1kg-Gwd is the Gambian dataset extracted from the 1kg-All dataset. The H3Africa dataset consists of the 1kg-Afr dataset in addition to other African datasets and can be accessed through the H3Africa imputation service. The HLA Multi-ethnic dataset (27)consists of datasets from the Japan Biological Informatics Consortium (28), the BioBank Japan Project (29), the Estonian Biobank (30), the 1kg-All (26)and a subset of studies in the TOPMed program (31).

The study set (Gambian dataset) was derived from the Gambian Genome Variation Project (GGVP) Whole Genome Sequence (WGS) dataset, a study that supports the discovery and understanding of genetic variants that influence human diseases. The population is made up of 4 different ethnic groups: Fula, Jola, Wolof and Mandinka. The GGVP project is a collaboration of the MRC Unit in the Gambia, the Wellcome Sanger Institute and the MRC Centre for Genomics and Global Health at Oxford University (32). The datasets are open access and can be found on the International Genome Sample Resource site (33). We chose the GGVP WGS dataset as the dataset was publicly available and worked with one target dataset as an example. Table 3 provides the sample size of each dataset.

**Table 3:**
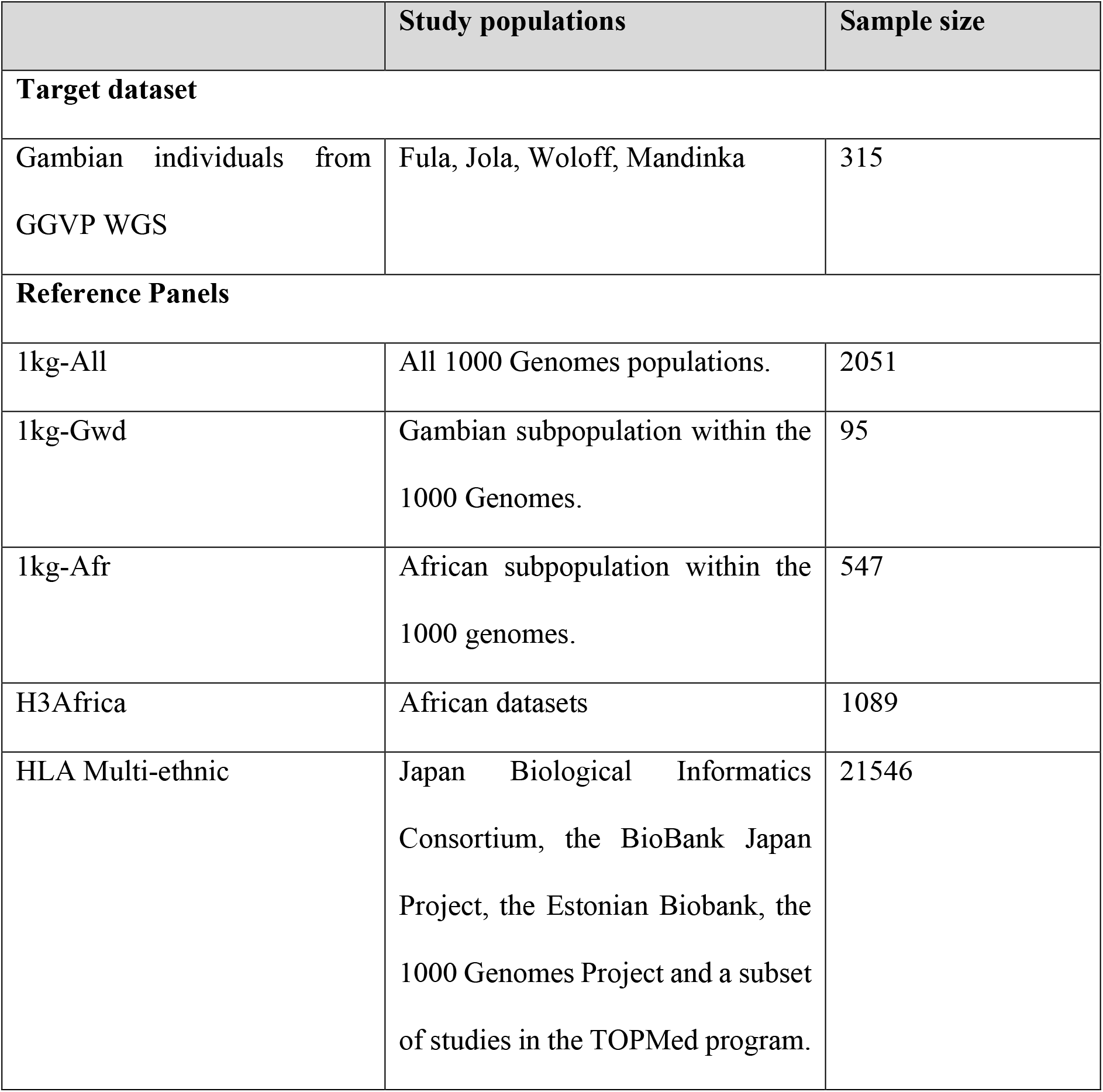
List of the study target datasets and reference panel populations.

To assess how the density of markers on the target dataset could affect the imputation performance of HLA alleles, we used two genotyping arrays commonly used for genomic studies of Africa populations, the Illumina Omni 2.5 array and the H3Africa array (34) with in-depth genomic coverage across diverse populations. The H3Africa array is based on the Illumina Omni 2.5 array, with approximately 75% markers overlapping with the Illumina Omni array, and the remaining 25% markers being custom-made. The Illumina Omni 2.5 array and the H3Africa array target datasets were created by selecting matching markers from the GGVP WGS datasets and masking the remaining SNPs.

### HLA imputation strategy

HLA imputation is the process of estimating a person’s HLA genotype from data on that person’s SNP genotypes at locations surrounding the HLA loci (35). The study focused on HLA class I alleles, which include HLA-A, HLA-B, and HLA-C alleles, because OptiType (36), the tool used to type true HLA alleles, only types class I HLA alleles.

Four imputation tools were used to impute HLA alleles. These included HLA allele specific imputation tools HIBAG version 1.4.0 with R statistical software version 3.6.1 (37), CookHLA (38), SNP2HLA (10), and a general imputation tool, Minimac4 (39). For SNP2HLA, PLINK version 1.07 was used for quality control, while BEAGLE version 3.0.4 was used for phasing and imputation.

HLA typing from WGS data was done using the OptiType (36) tool in the nf-core HLA typing pipeline (40,41). Then, using python scripts, HLA types were combined following the required format for HIBAG, CookHLA and SNP2HLA. SNP genotypes were also converted to PLINK format using PLINK version 2.0.

For the reference panel, one can use a ready-made reference panel or create a custom reference panel using HLA types and SNP genotypes. For this analysis, one reference panel (HLA Multi-ethnic) (27) was ready-made while four reference panels (1kg-All, 1kg-Afr, 1kg-Gwd, H3Africa) were custom-made.

For HIBAG, SNP2HLA and CookHLA, custom-made reference panels were constructed using HLA types and SNP genotypes. A genetic map was generated using the ‘MakeGeneticMap’ module within the CookHLA package prior to training the reference panel. We used the “MakeReference” module in SNP2HLA to construct the SNP2HLA and CookHLA reference panels, and the “hlaAttrBagging” function in HIBAG to train the HIBAG specific reference panels. For Minimac4, reference panels were generated using SNP genotypes and HLA alleles typed using the HLA-LA tool (42) instead of OptiType (36). This was to match the method used to create the HLA Multi-ethnic reference panel and thus enable comparison.

HLA alleles were then imputed from SNP data using the ‘SNP2HLA’ script with window size set to the default of 1000 for SNP2HLA and the hlaPredict function for HIBAG. For CookHLA, the ‘CookHLA.py’ script was used for imputation. For Minimac4, HLA alleles were imputed by calling the Minimac4 tool. For the HLA Multi-ethnic reference panel, the sample datasets were submitted to the Michigan imputation server (43) and HLA imputation was conducted using the Minimac4 imputation tool.

### Imputation accuracy assessment

We used concordance rate as the primary assessment metric, which is the percentage of correctly imputed best-guess alleles of all imputed alleles based on true HLA alleles. The true HLA alleles were obtained by typing HLA alleles using OptiType tool that has been shown to type HLA Class I allele at 99% accuracy (44). The hlaCompareAllele function in HIBAG was used to calculate the concordance rate, while the ‘measureacc’ module in the CookHLA package (38) was used to calculate the SNP2HLA, CookHLA and Minimac4 concordance rate.

Another interesting way of looking at the accuracy results is by computing HLA allele error rates, which is done by subtracting the accuracy values from 1. Allele frequencies reflect the genetic diversity in a population. As HLA alleles are more genetically diverse, computing HLA allele frequency is important in establishing the accuracy of HLA alleles. HLA allele frequencies were computed using the PyPop (45) package and a comparison was made between the allele frequencies and concordance rates of HLA alleles.

### Reproducibility

To enhance reproducibility, GitHub was used for documentation and version control and the tools were packaged and deployed using Docker and Singularity containers. Containers are a lightweight form of virtualization, used to package and distribute applications. The packaged tools include htslib, Samtools, VCFtools, Minimac4, Bcftools, Bedtools, SNP2HLA, CookHLA, HIBAG, Plink2, Miniconda3, R, Java, Perl, and R Libraries. In addition, the Nextflow (46) workflow language was used to automate the pipeline and the tools packaged using Docker and Singularity containers. The documentation is available on GitHub (47).

A summary of the workflow used for the analysis is presented in Fig 4 below. Matching markers from the GGVP WGS datasets were chosen to produce the target datasets for the Illumina Omni 2.5 and the H3Africa arrays. The datasets were then imputed on 5 reference panels using 4 imputation tools, and HLA imputation accuracy was assessed using concordance rate.

**Fig 4:**
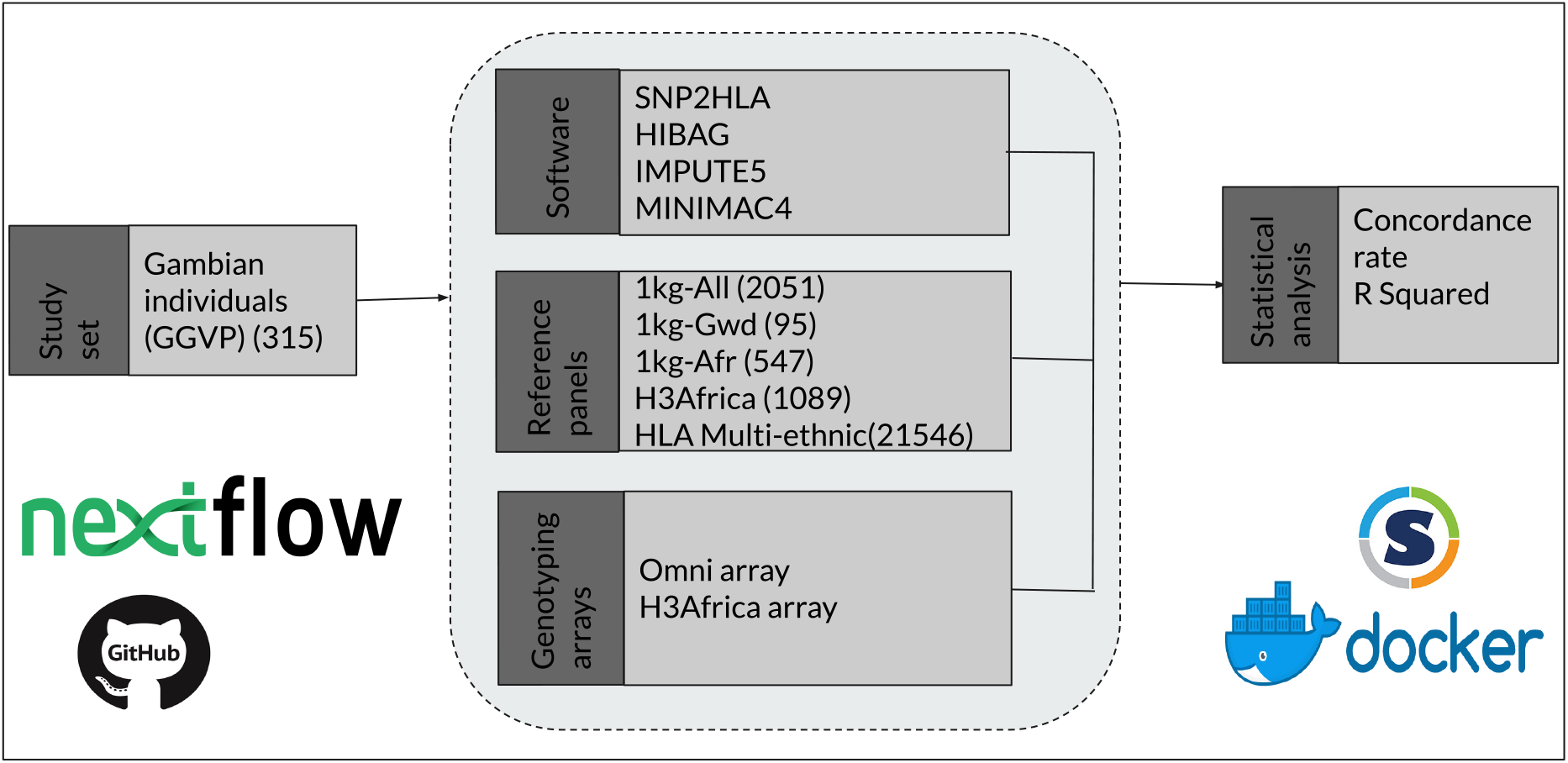
A summary of the materials and methods used.

## Ethics statement

Ethical approval was not required as the study was benchmarking imputation tools and reference panels using publicly available datasets (1000 Genomes and GGVP projects). Additional reference panels used are accessible through a public imputation service.

## Code availability

The pipeline can be accessed at https://github.com/nanjalaruth/MHC-Imputation-Accuracy

## Data availability

The 1000 genomes and GGVP WGS datasets are open source and can be accessed from the International Genome Sample Resource. The H3Africa reference panel can be accessed for imputation by requesting access to run datasets via the imputation service.

## Acknowledgments

The Computational Biology Division within the University of Cape Town, South Africa for providing access to the Ilifu cloud server.

## References

1. Cao H, Wu J, Wang Y, Jiang H, Zhang T, Liu X, et al. An Integrated Tool to Study MHC Region: Accurate SNV Detection and HLA Genes Typing in Human MHC Region Using Targeted High-Throughput Sequencing. PLoS One [Internet]. 2013 Jul 24 [cited 2022 Feb 10];8(7): e69388. Available from: https://journals.plos.org/plosone/article?id=10.1371/journal.pone.0069388

2. Degenhardt F, Wendorff M, Wittig M, Ellinghaus E, Datta LW, Schembri J, et al. Construction and benchmarking of a Multi-ethnic reference panel for the imputation of HLA class I and II alleles. Hum Mol Genet. 2019;28(12):20782092.

3. Huang Y, Yang J, Ying D, Zhang Y, Shotelersuk V, Hirankarn N, et al. HLAreporter: a tool for HLA typing from next generation sequencing data. Genome Med [Internet]. 2015 Mar 16 [cited 2022 Dec 21];7(1). Available from: https://pubmed.ncbi.nlm.nih.gov/25908942/

4. Shiina T, Hosomichi K, Inoko H, Kulski JK. The HLA genomic loci map: Expression, interaction, diversity and disease. Vol. 54, Journal of Human Genetics. 2009. p. 15–39.

5. Khor SS, Yang W, Kawashima M, Kamitsuji S, Zheng X, Nishida N, et al. High-Accuracy imputation for HLA class i and II genes based on high-resolution SNP data of population-specific references. Pharmacogenomics Journal [Internet]. 2015;15(6):530–7. Available from: http://dx.doi.org/10.1038/tpj.2015.4

6. Dendrou CA, Petersen J, Rossjohn J, Fugger L. HLA variation and disease. Nat Rev Immunol [Internet]. 2018;18(5):325–39. Available from: http://dx.doi.org/10.1038/nri.2017.143

7. Naj AC. Genotype Imputation in Genome-Wide Association Studies. Curr Protoc Hum Genet. 2019;102(1):1–15.

8. Sherry ST, Ward MH, Kholodov M, Baker J, Phan L, Smigielski EM, et al. DbSNP: The NCBI database of genetic variation. Nucleic Acids Res. 2001;29(1):308–11.

9. Choo SY. The HLA System: Genetics, Immunology, Clinical Testing, and Clinical Implications. Yonsei Med J [Internet]. 2007 Feb [cited 2022 Jan 27];48(1):11. Available from: /pmc/articles/PMC2628004/

10. Jia X, Han B, Onengut-Gumuscu S, Chen WM, Concannon PJ, Rich SS, et al. Imputing Amino Acid Polymorphisms in Human Leukocyte Antigens. PLoS One [Internet]. 2013 Jun 6 [cited 2022 Feb 10];8(6):e64683. Available from: https://journals.plos.org/plosone/article?id=10.1371/journal.pone.0064683

11. Lonjou C, Zhang W, Collins A, Tapper WJ, Elahi E, Maniatis N, et al. Linkage disequilibrium in human populations. Proc Natl Acad Sci U S A [Internet]. 2003 May 5 [cited 2022 Dec 2];100(10):6069. Available from: /pmc/articles/PMC156327/

12. Choudhury A, Aron S, Botigué LR, Sengupta D, Botha G, Bensellak T, et al. High-depth African genomes inform human migration and health. Nature. 2020;586(7831):741–8.

13. Gurdasani D, Carstensen T, Tekola-Ayele F, Pagani L, Tachmazidou I, Hatzikotoulas K, et al. The African Genome Variation Project shapes medical genetics in Africa. Nature 2014 517:7534 [Internet]. 2014 Dec 3 [cited 2022 Nov 22];517(7534):327–32. Available from: https://www.nature.com/articles/nature13997

14. Gronau I, Hubisz MJ, Gulko B, Danko CG, Siepel A. Bayesian inference of ancient human demography from individual genome sequences. Nat Genet. 2011;43(10):1031–5.

15. Nielsen R, Akey JM, Jakobsson M, Pritchard JK, Tishkoff S, Willerslev E. Tracing the peopling of the world through genomics. Nature. 2017;541(7637):302–10.

16. Gross M. African genomes. Current Biology [Internet]. 2011;21(13):R481–4. Available from: http://dx.doi.org/10.1016/j.cub.2011.06.047

17. Schurz H, Müller SJ, van Helden PD, Tromp G, Hoal EG, Kinnear CJ, et al. Evaluating the accuracy of imputation methods in a five-way admixed population. Front Genet. 2019;10(february):1–9.

18. Levin AM, Adrianto I, Datta I, Iannuzzi MC, Trudeau S, McKeigue P, et al. Performance of HLA allele prediction methods in African Americans for class II genes HLA-DRB1, −DQB1, and –DPB1. BMC Genet [Internet]. 2014 Jun 16 [cited 2022 Nov 27];15:72. Available from: /pmc/articles/PMC4074844/

19. Karnes JH, Shaffer CM, Bastarache L, Gaudieri S, Glazer AM, Steiner HE, et al. Comparison of HLA allelic imputation programs. PLoS One. 2017;12(2):1–12.

20. Moutsianas L, Gutierrez-Achury J. Genetic association in the HLA region. In: Methods in Molecular Biology. 2018. p. 111–34.

21. Raghavan M, Geng J. HLA-B polymorphisms and intracellular assembly modes. Mol Immunol. 2015;68(2):89–93.

22. Robinson J, Halliwell JA, Hayhurst JD, Flicek P, Parham P, Marsh SGE. The IPD and IMGT/HLA database: Allele variant databases. Nucleic Acids Res. 2015;43(D1): D423–31.

23. Carrington M, Walker BD. Immunogenetics of spontaneous control of HIV. Vol. 63, Annual Review of Medicine. 2012. p. 131–45.

24. Verlouw JAM, Clemens E, de Vries JH, Zolk O, Verkerk Ajmh, am Zehnhoff-Dinnesen A, et al. A comparison of genotyping arrays. European Journal of Human Genetics [Internet]. 2021;29(11):1611–24. Available from: https://doi.org/10.1038/s41431-021-00917-7

25. Browning BL, Browning SR. A unified approach to genotype imputation and haplotype-phase inference for large data sets of trios and unrelated individuals. Am J Hum Genet [Internet]. 2008;84(2):210–23. Available from: http://dx.doi.org/10.1016/j.ajhg.2009.01.005

26. Auton A, Abecasis GR, Altshuler DM, Durbin RM, Bentley DR, Chakravarti A, et al. A global reference for human genetic variation. Nature [Internet]. 2015 Sep 30 [cited 2022 Nov 22];526(7571):68–74. Available from: https://pubmed.ncbi.nlm.nih.gov/26432245/

27. Luo Y, Kanai M, Choi W, Li X, Sakaue S, Yamamoto K, et al. A high-resolution HLA reference panel capturing global population diversity enables multi-ancestry fine-mapping in HIV host response. Nat Genet [Internet]. 2021 Oct 1 [cited 2022 Nov 2];53(10):1504–16. Available from: https://pubmed.ncbi.nlm.nih.gov/34611364/

28. Hirata J, Hosomichi K, Sakaue S, Kanai M, Nakaoka H, Ishigaki K, et al. Genetic and phenotypic landscape of the major histocompatibilty complex region in the Japanese population. Nat Genet [Internet]. 2019 Mar 1 [cited 2022 Nov 22];51(3):470–80. Available from: https://pubmed.ncbi.nlm.nih.gov/30692682/

29. Okada Y, Momozawa Y, Sakaue S, Kanai M, Ishigaki K, Akiyama M, et al. Deep whole-genome sequencing reveals recent selection signatures linked to evolution and disease risk of Japanese. Nature Communications 2018 9:1 [Internet]. 2018 Apr 24 [cited 2022 Nov 22];9(1):1–10. Available from: https://www.nature.com/articles/s41467-018-03274-0

30. Nelis M, Esko T, Mägi R, Zimprich F, Toncheva D, Karachanak S, et al. Genetic Structure of Europeans: A View from the North–East. PLoS One [Internet]. 2009 May 8 [cited 2022 Nov 22];4(5):e5472. Available from: https://journals.plos.org/plosone/article?id=10.1371/journal.pone.0005472

31. NHLBI Trans-Omics for Precision Medicine WGS-About TOPMed [Internet]. [cited 2022 Nov 22]. Available from: https://topmed.nhlbi.nih.gov/

32. GGVP GRCh38 | IGSR data collection [Internet]. [cited 2022 Aug 17]. Available from: https://www.internationalgenome.org/data-portal/data-collection/ggvp-grch38

33. Fairley S, Lowy-Gallego E, Perry E, Flicek P. The International Genome Sample Resource (IGSR) collection of open human genomic variation resources. Nucleic Acids Res. 2020 Jan 1;48(D1):D941–7.

34. H3Africa array annotations [Internet]. [cited 2022 Nov 22]. Available from: https://chipinfo.h3abionet.org/

35. Meyer D, Nunes K. HLA imputation, what is it good for? Hum Immunol. 2017 Mar 1;78(3):239–41.

36. Szolek A, Schubert B, Mohr C, Sturm M, Feldhahn M, Kohlbacher O. OptiType: precision HLA typing from next-generation sequencing data. Bioinformatics [Internet]. 2014 Dec 1 [cited 2022 Sep 5];30(23):3310–6. Available from: https://pubmed.ncbi.nlm.nih.gov/25143287/

37. Zheng X, Shen J, Cox C, Wakefield JC, Ehm MG, Nelson MR, et al. HIBAG - HLA genotype imputation with attribute bagging. Pharmacogenomics Journal [Internet]. 2014;14(2):192–200. Available from: http://dx.doi.org/10.1038/tpj.2013.18

38. Luo Y, Cook S, Choi W, Lim H, Kim K, Jia X, et al. Accurate imputation of human leukocyte antigens with CookHLA. Nat Commun [Internet]. 2021;12(1):1–11. Available from: http://dx.doi.org/10.1038/s41467-021-21541-5

39. van Leeuwen EM, Kanterakis A, Deelen P, Kattenberg M v., Slagboom PE, de Bakker Piw, et al. Population-specific genotype imputations using minimac or IMPUTE2. Nat Protoc [Internet]. 2015;10(9):1285–96. Available from: http://dx.doi.org/10.1038/nprot.2015.077

40. nf-core/hlatyping: Precision HLA typing from next-generation sequencing data [Internet]. [cited 2022 Nov 22]. Available from: https://github.com/nf-core/hlatyping

41. Ewels PA, Peltzer A, Fillinger S, Patel H, Alneberg J, Wilm A, et al. The nf-core framework for community-curated bioinformatics pipelines. Nat Biotechnol. 2020 Mar 1;38(3):276–8.

42. Dilthey AT, Mentzer AJ, Carapito R, Cutland C, Cereb N, Madhi SA, et al. HLA*LA—HLA typing from linearly projected graph alignments. Bioinformatics [Internet]. 2019 Nov 1 [cited 2022 Aug 27];35(21):4394–6. Available from: https://academic.oup.com/bioinformatics/article/35/21/4394/5426702

43. Michigan Imputation Server [Internet]. [cited 2022 Nov 22]. Available from: https://imputationserver.sph.umich.edu/index.html#!

44. Yi J, Chen L, Xiao Y, Zhao Z, Su X. Investigations of sequencing data and sample type on HLA class Ia typing with different computational tools. Brief Bioinform [Internet]. 2021 May 20 [cited 2022 Sep 5];22(3):1–6. Available from: https://academic.oup.com/bib/article/22/3/bbaa143/5871189

45. (8) (PDF) PyPop User Guide: User Guide for Python for Population Genomics [Internet]. [cited 2022 Sep 6]. Available from: https://www.researchgate.net/publication/271852987_PyPop_User_Guide_User_Guide_for_Python_for_Population_Genomics

46. A DSL for parallel and scalable computational pipelines | Nextflow [Internet]. [cited 2022 Nov 22]. Available from: https://www.nextflow.io/

47. nanjalaruth/MHC-Imputation-Accuracy: A project on evaluating the accuracy of genotype imputation in the human MHC region in selected African populations. [Internet]. [cited 2022 Nov 22]. Available from: https://github.com/nanjalaruth/MHC-Imputation-Accuracy

